# Differential effects of sodium on agonist-induced conformational transitions and signaling at μ and κ opioid receptors

**DOI:** 10.1101/2025.08.28.672897

**Authors:** Bram Kuijer, Camryn J. Fulton, Talia L. Albert, Paloma M. Knobloch, Jing Wang, Brian E. Krumm, Tao Che, Balazs R. Varga, Susruta Majumdar, Vsevolod Katritch, Ryan H. Gumpper, Terry Kenakin, Xi-Ping Huang, Bryan L. Roth

**Author notes:** =Equal contributions (Bram Kuijer, Camryn J. Fulton).

## Abstract

Sodium ions are classically conceptualized as negative allosteric modulators for G protein coupled receptors (GPCRs), although there have been reports of either positive allosteric modulation or no effect of sodium on GPCR function. Here we identified opposing actions of sodium on the μ and κ opioid receptors. We utilized a variety of methods including radioligand binding, real-time conformational monitoring of transitions using bioluminescence resonance energy transfer and signaling assays using the TRUPATH resource. At the μ receptors, sodium behaved as a negative allosteric modulator of binding, conformational transitions and signaling. Intriguingly, bitopic μ agonists were unaffected by sodium concentrations. By contrast, at the κ opioid receptor sodium negatively modulated agonist binding and positively modulated conformational transitions and signaling. Taken together, these findings support the notion that the differential sensitivities to sodium concentrations will result in opposing effects on cell surface and intracellular signaling.

## INTRODUCTION

G protein-coupled receptors (GPCRs) represent both the single largest class of mammalian transmembrane receptors and the molecular targets for about 1/3^rd^ of FDA-approved medications (*1*). Among the various GPCRs, opioid receptors are of interest because they mediate the actions of many analgesics and drugs of abuse. The μ opioid receptor (MOR), for example, is essential for the actions of analgesic drugs like morphine, (*2*) while the κ opioid receptor (KOR) is the target of psychotomimetic compounds like salvinorin A (*3*).

The opioid receptors were initially discovered in 1973 (*4–6*) via radioligand binding technology and are divided into four main classes: μ, 8, κ, and nociception (*7*). Shortly after their discovery, opioid receptors were shown to be allosterically regulated by physiological concentrations of Na^+^ (*8, 9*) (*10*). Over the past several years, the structures of the inactive and active states of opioid receptors have been solved, providing insights into the molecular details of actions of these drugs (*11–18*). A high-resolution crystal structure of the δ opioid receptor (DOR) revealed that the Na^+^ site is distinct from the orthosteric site occupied by opioid agonists and antagonists (*19*). This site is predicted to be conserved among most GPCRs including all opioid receptors (*20*) even though the sodium site has only been directly visualized in a handful of GPCRs (see (*21–23*)) as, typically, resolutions < 2Å are required. Despite the lack of direct structural details, a large number of papers have demonstrated sodium’s allosteric action in promoting inactive, transducer uncoupled states for many GPCRs (*24*) (*25*). Recent studies have taken advantage of the sodium site as part of a bitopic ligand design strategy at the μ-opioid, δ-opioid, and CB₁ cannabinoid receptors. Notably, the CB₁ cannabinoid receptor is sodium insensitive.

Recently, we identified two opioid-directed nanobodies –Nb39 (*13, 15*) and Nb6 (*14, 15*)—which stabilize the active (Nb39) and inactive (Nb6) states of κ (KOR) and μ (MOR) opioid receptors, respectively. As we demonstrated by utilizing a bioluminescence resonance energy transfer (BRET) approach, we could quantify agonist- and antagonist-specific conformational transitions in real time (*14*) using nanobody-based BRET-sensors.

Utilizing this dual nanobody-based biosensor approach, radioligand binding, and our TRUPATH platform (*26*) to monitor the activation of heterotrimeric G proteins, we found that sodium differed in the ability to modulate agonist and antagonist conformational transitions at MOR and KOR. At MOR, sodium behaved as a negative allosteric modulator (NAM), while at KOR, sodium enhanced agonist potency and efficacy while negatively modulating radioligand binding. Interestingly, the NTSR1 neurotensin receptor, used as a control for the signaling assays, was relatively insensitive to sodium. These results indicate that sodium’s allosteric actions are complex and are dependent upon the receptor and its transducer.

## EXPERIMENTAL DETAILS

### Chemistry

C5 guano and C6 guano were synthesized as previously described(*27*).

### Radioligand binding assays and data analysis

Radioligand binding assays were carried out using human MOR or KOR opioid membrane preparations made from transiently transfected Expi293 cells in 96-well plates at the final volume of 125 µul per well. Briefly, For KOR binding assays, ^3^H-U69593 serial dilutions were incubated with KOR membranes in the absence and presence of 140 mM NaCl at the room temperature in the dark for 60 minutes. Salvinorin A (Sal A, 10 µM final) was included to define nonspecific binding. For MOR binding assays, receptor membranes and [^3^H]-Naloxone (around K_d_ levels) were incubated with 12-point serial dilutions of MOR agonist DAMGO or antagonist naltrexone in the absence and presence of fixed concentrations of NaCl for 90 min at room temperature in the dark. To reach the final concentrations of NaCl, DAMGO or Naltrexone serial dilutions were prepared in the binding buffer (50 mM Tris HCl, 10 mM MgCl2, 0.1 mM EDTA, pH 7.40) at the 5x of the final concentrations and added to 96-well plate at 25 µl/well. The reactions were stopped by vacuum filtration onto 96-well UniFilter GF/C filter plates, briefly socked in the cold 0.3% (w/v) polyethyleneimine solution just before use. The Filter plates were oven-dried before receiving Microscint-O scintillation cocktail. Radioactivity was counted using a Wallac MicroBeta TriLux counter. Results were normalized (100 and 0% corresponding to the binding in the absence and presence of competing ligand under control condition – no Sodium ions) and analyzed using the GraphPad Prism V10 (detail below). KOR saturation binding results were fitted to the built-in saturation function (one-site total and nonspecific) in the Prism.

To obtain allosteric parameters, the MOR competition binding results were fitted to the following allosteric binding function based on the simplified allosteric binding model (Fig 1A), in which an agonist (DAMGO) or an antagonist (Naltrexone) competes with radioligand ^3^H-Naloxone in the absence and presence of specified concentration of sodium.

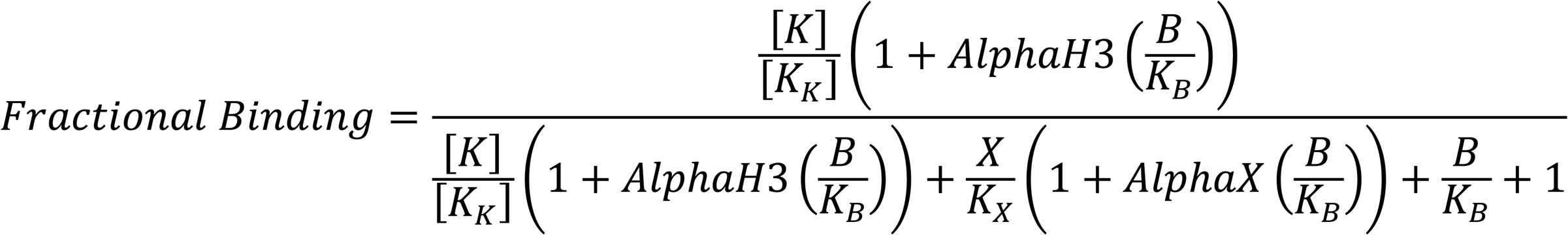

**Fig 1.**
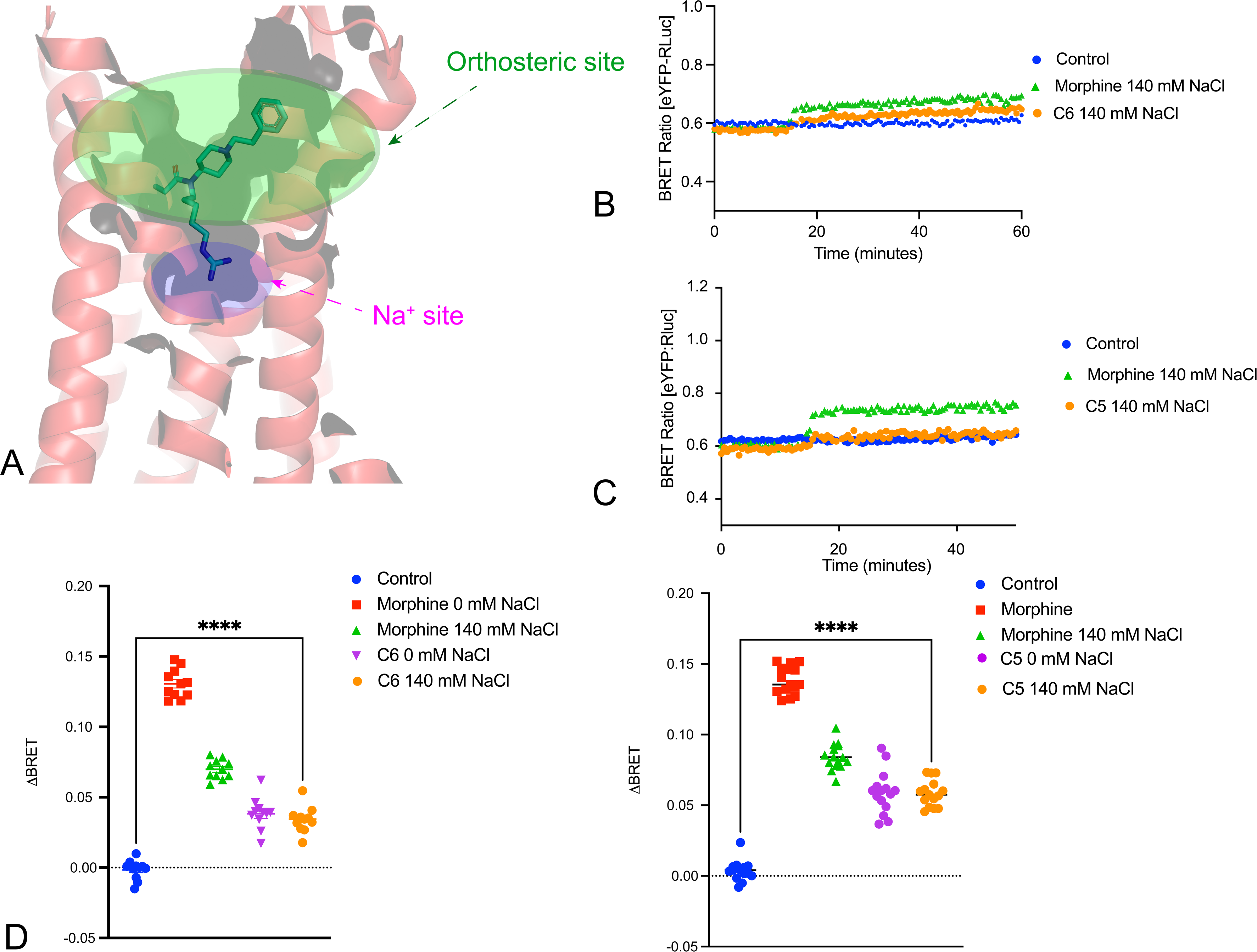
Allosteric binding in the absence and presence of NaCl. TOP, allosteric binding model showing a receptor (R) allowing the concomitant binding of sodium (Na), a radioligand (L), and a competing agonist or antagonist (M), with corresponding binding constants. In this case, an agonist (Morphine or DAMGO for MOR) or an antagonist (Naltrexone) binds to ^3^H-Naloxone labeled MOR in the presence of NaCl. NaCl has independent allosteric modulation on Naloxone and competing ligand. A and C, DAMGO and Naltrexone binding curves. B and D, overlay 3D views of actual DAMGO and Naltrexone binding points (dots) on top of simulated results with parameters from panels A and C. Simulations were performed in the GraphPad Prism V10 and the 3D views were plotted with the SigmaPlot V16.

It is assumed that sodium ions have independent allosteric modulations on the radioligand [^3^H]-Naloxone (AlphaH3) and competing orthosteric agonist DAMGO or antagonist Naloxone (AlphaX). In the model, K is the radioligand concentration, usually at its K_d_ level. B is the sodium concentration. K_B_ and K_X_ are binding affinity of allosteric modulator Sodium and competing ligand. AlphaH3, AlphaX, K_B_, and K_X_ are the parameters to be resolved by global fitting to sets of data with shared parameters in the absence and presence of Sodium ions. Alpha > 1 for positive modulation and 0 < alpha < 1 for negative modulation.

### TRUPATH BRET2 Protocol

Cells were trypsinized and plated in 10 cm dishes at 7-8 x 10^6^ cells/dish or in 6 cm dishes at a density of 2.5 x 10^6^ cells/dish in DMEM complete medium (500 ml DMEM, Genesee cat no 25-500; 50 ml FBS, Gen Clone cat no 25-514; 5 ml Pen/Strep, Gen Clone cat no 25-512). After 5-6 hours, cells were transfected with 1μg of plasmid for 10cm dishes or 250ng of plasmid for 6 cm dishes in a 1:1:1:1 mixture of receptor:⍺:β:ɣ, and the recommended 3 *μ* L/μg of Transit-2020 (Mirus cat. no. MIR5406) in 200 μL of Opti-MEM (Gibco cat. no. 31985-070). The transfection mixture was incubated for 45 minutes before being added dropwise to the dish. The following day, the media was aspirated and replaced with DMEM containing 1% dialyzed FBS. After resting for 3-4 hours, cells were removed from the dishes using Versene (1X PBS, 0.5 mM EDTA, pH 7.4), and plated in Corning 96-well flat clear bottom white microplates at a density of 30,000-50,000 cells/well in 200 μL of DMEM with 1% dialyzed FBS (dFBS). After overnight incubation, medium was aspirated from the plate and replaced with 60 μL of assay buffer (20 mM HEPES, 1.26 mM calcium chloride, 5.33 mM potassium chloride, 0.44 mM potassium phosphate (monobasic), 0.30 mM potassium phosphate (dibasic), 0.5 mM magnesium chloride, 0.41 mM magnesium sulfate, and 5.60 mM glucose) containing either 140 mM NaCl or 140 mM Choline Cholide (ChCl). Dilutions were made in drug buffer (140 mM NaCl or ChCl assay buffer plus 0.3% Bovine Serum Albumin and 3 mM Ascorbic acid). After a 10-minute incubation at 37℃, 30 μL of drug dilutions were added to each well and incubated for another 10 minutes at room temperature. Following this incubation, 10 µL Coelenterazine 400a (CZT-400a, Nanolight, cat. no. 340-500) was added to each well, giving a final concentration of 5 µM. Plates were incubated for 10 minutes in the dark before reading. Plates were read using PHERAstar FSX (BMG LabTech, software v5.41).

### Kinetic studies

HEK293T cells were cultured in-house and initially grown in 15 cm dishes prior to experimental use. Cells were trypsinized and seeded in 6-well plates at a density of 7.25 × 10⁵ cells/well in 2 mL DMEM complete medium as above. After 5 hours, cells were transfected with a 100 µL transfection mixture per well, consisting of MOR-RLuc (200 ng), Nb6 or Nb39 (500 ng), Opti-MEM (Gibco cat. no. 31985-070), and TransIT-2020 (3 µL/µg DNA, Mirus cat. no. MIR5406). Transfection mix was prepared at room temperature and incubated for 45 minutes in the dark before addition. All HEK293T cells and plasmids used for transfection were generated in-house. Each experiment was conducted in biological triplicate (n = 3).

The following day, cells were trypsinized and replated into Greiner 96-well F-bottom plates (cat. no. 655098) at a concentration of 35,500 cells/well in DMEM with 1% dFBS (500 mL DMEM, 5 mL dFBS, 5 mL Pen/Strep). On day three, medium was aspirated, light-blocking tape was applied to the plate bottoms, and cells were washed with Na⁺-free assay buffer.

Cells were incubated for 10 minutes at 37°C with 60 µL of either 0 mM, 60 mM, or 140 mM NaCl (with choline chloride added to maintain osmolarity). Following incubation, 10 µL Coelenterazine h (Promega, cat. no. S201A) was added to a final concentration of 5 µM and incubated in the dark at room temperature. Plates were then read using a PHERAstar FSX multi-mode reader (BMG LabTech, software v5.41) configured for BRET1 kinetic assays: 30-minute cycle time, 0.16 s measurement interval, simultaneous dual emission, gain = 3000 for channels A and B. Readings were taken for 30 baseline cycles. Then, 30 µL of agonist (C5, C6, DAMGO, or morphine) was added per well in triplicate at varying NaCl concentrations. At cycle 130, 10 µL naloxone (final well concentration = 10 µM) was added; kinetic readings continued through cycle 230.

### Data Analysis

All data were analyzed using GraphPad Prism 10.0 (GraphPad Software, San Diego, CA). Dose-response curves were fitted using a three-parameter nonlinear regression model (log[agonist] vs. response). EC₅₀ values and span were derived directly from curve fits. Data are represented as means ± SEM from three independent experiments (n = 3).

## RESULTS

### Modeling sodium’s allosteric actions on opioid receptor binding

Many prior studies have shown that sodium allosterically decreases agonist binding affinity and enhances antagonist binding at opioid receptors (*8*) (*10*). High-resolution structural studies have identified a preformed sodium pocket separate from the orthosteric site on opioid receptors (*23*) and other GPCRs (*21*) (*28*) (*24*). As shown in Fig 1, sodium is predicted to have a complex action on opioid receptor binding properties, displaying positive allosteric modulation of antagonist binding and negative allosteric modulation of agonist binding. To quantify the various allosteric parameters describing the effect of sodium on agonist and antagonist potencies, a more precise approach to analyzing the binding data was developed by first creating a thermodynamic model describing the independent interaction of sodium as an allosteric modulator. We also considered that antagonists and agonists compete for the same orthosteric site. As also shown in Fig 1, the allosteric equation and fitted parameters (Table 1) were able to fit experimental data well. These findings are consistent with a model in which sodium functions as a NAM for agonists and a positive allosteric modulator (PAM) for antagonists.

**Table 1.** Allosteric parameters. Competition binding results were normalized to control conditions in the absence of NaCl, where 100% and 0% for the binding in the absence and presence of competing ligand at the highest concentration. Pooled results were fitted to the allosteric binding model (see Methods) with an average ^3^H-Naloxone concentration ([K]) at 1.38 nM and average binding affinity (K_K_) at 3.31 nM. The binding affinity of competing ligands (K_X_) in the absence of NaCl was determined by fitting the results with the built-in One-site K_i_ function in Prism and constrained. They are 22.91 nM, 28.84 nM, 23.44 nM, and 2.88 nM for DAMGO, Morphine, Naltrindole, and Naltrexone, respectively. All constrained parameters were shared in fitting for each date set.

| pK<sub>B</sub> (Sodium binding affinity)<br>(K<sub>B</sub> in mM) | LogAlphaH3<br>(AlphaH3) | LogAlphaX<br>(AlphaX) |
| --- | --- | --- |
| DAMGO competition binding to [$^3\text{H}$]-Naloxone labeled MOR ± NaCl | | |
| 2.66 ± 0.03<br>(2.19 mM) | 0.17 ± 0.01<br>(1.48) | -2.22 ± 0.10<br>(0.0062) |
| Morphine competition binding to [$^3\text{H}$]-Naloxone labeled MOR ± NaCl | | |
| 2.67 ± 0.03<br>(2.14 mM) | 0.19 ± 0.01<br>(1.54) | -1.22 ± 0.03<br>(0.059) |
| Naltrindole competition binding to [$^3\text{H}$]-Naloxone labeled MOR ± NaCl | | |
| 1.91 ± 0.05<br>(123 mM) | 0.43 ± 0.01<br>(2.70) | 0.59 ± 0.02<br>(3.89) |
| Naltrexone competition binding to [$^3\text{H}$]-Naloxone labeled MOR ± NaCl | | |
| 2.43 ± 0.06<br>(3.73 mM) | 0.25 ± 0.01<br>(1.76) | -0.37 ± 0.03<br>(0.42) |

### Real-time visualization of sodium-stabilized conformational transitions at MOR

Given this model for describing sodium’s actions, we next tested predictions that one could make using this type of model in real time in live cells *in situ*. For these studies, we evaluated the ability of nanobodies to report conformational transitions at MOR with HEK293T cells co-transfected with MOR-RLuc and Nb6-mVenus or Nb39-mVenus (*14, 29–31*). BRET ratios were quantified in real-time (*14, 15*) following agonist (morphine; 10 μM final concentration) or antagonist (naloxone; 10 μM final concentration) exposure. Nb6 has been shown to bind to the inactive-state and, upon agonist addition, dissociates from the receptor. As shown in Fig 2A, morphine caused a rapid decline in the MOR-Rluc/Nb6-mVenus BRET response. This response is consistent with a morphine-stabilized active-state conformation leading to dissociation of Nb6. In a similar manner, we evaluated Nb39 as a probe for MOR-dependent active state conformational transitions. As shown in Fig 3B, morphine (10 μM) rapidly promoted an increase in the BRET ratio, indicating Nb39 recruitment to the receptor and stabilization of the active-state.

**Fig 2.**
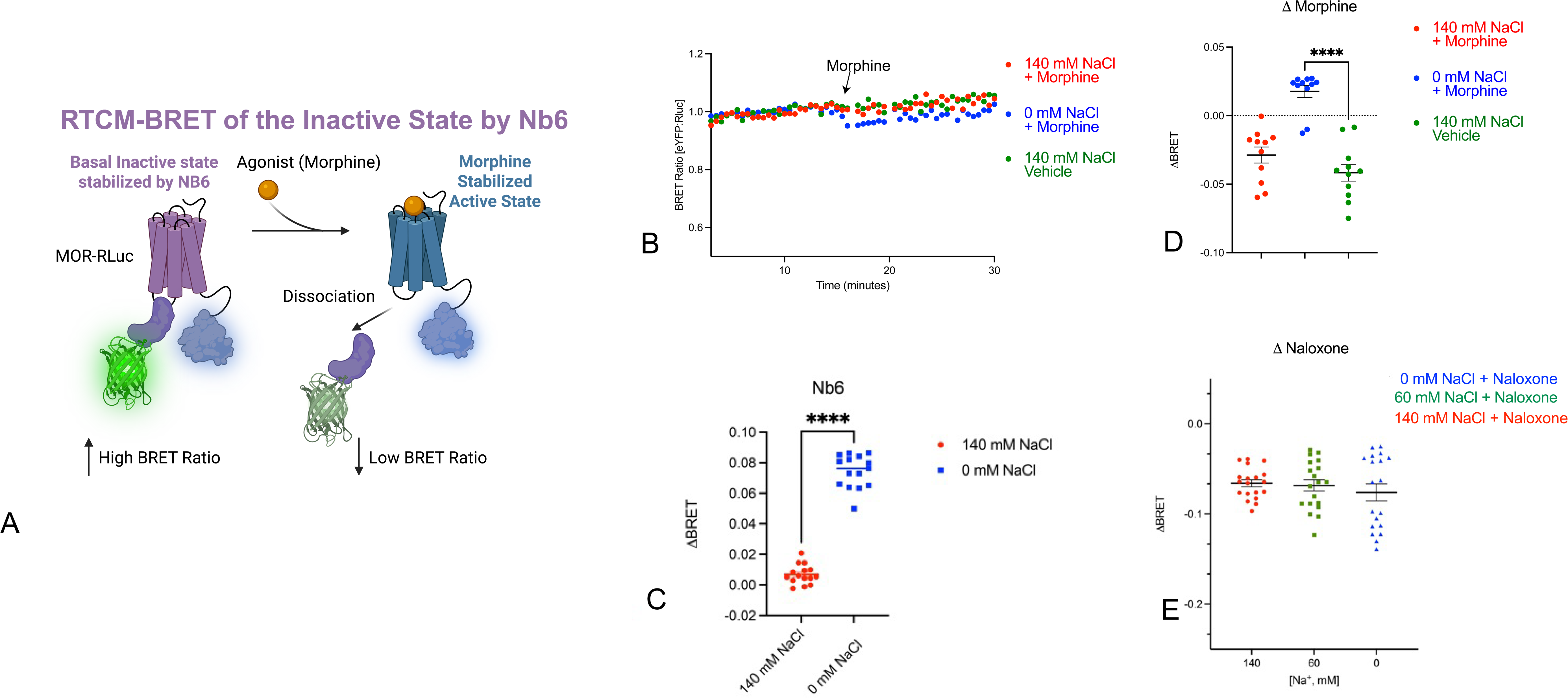
Real-Time Conformational Measurement (RTCM) BRET at MOR using the inactive-state Nb6 nanobody. Panel A shows in cartoon form the overall approach whereby agonist binding leads to an active state which has low affinity for Nb6, thereby reducing the BRET signal, B shows a typical time course at two different sodium concentrations. C shows the difference between the basal BRET at two different sodium concentrations averaged from multiple time points with 3 independent biological replicates. D shows the response due to the agonist morphine for N=3 independent biological replicates; E shows the response due to the antagonist naloxone at 3 different sodium concentrations for N=3 independent biological replicates. For all: ****p<0.001

**Fig 3.**
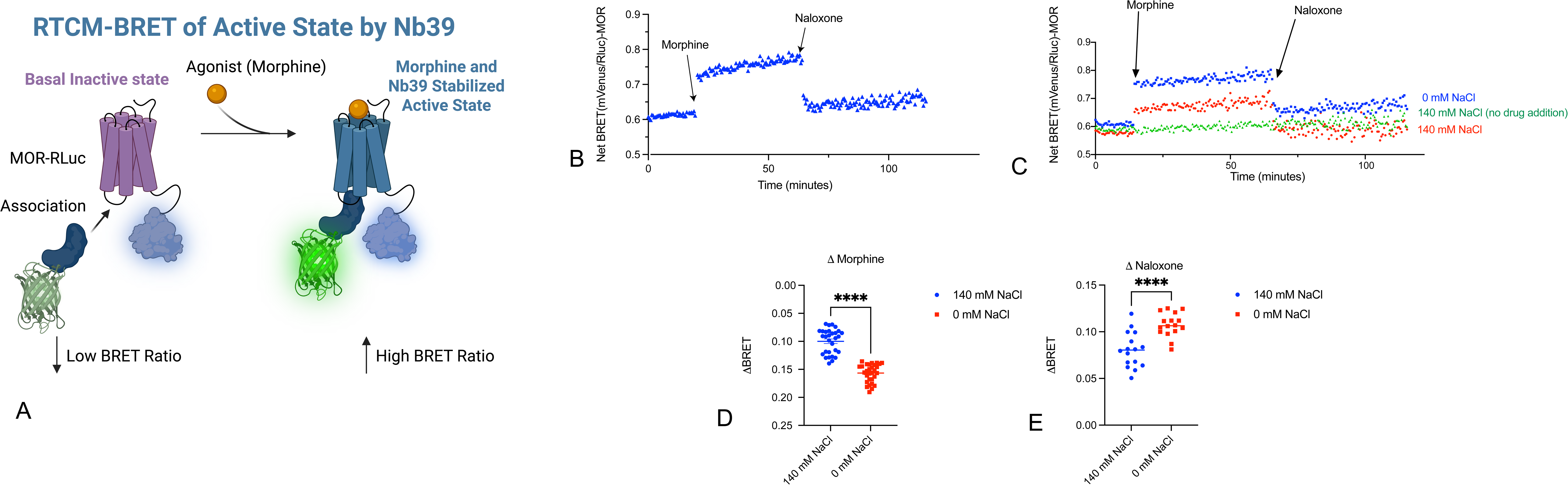
Real-Time Conformational Measurement (RTCM) BRET at MOR using the active-state Nb39 nanobody. Panel A shows in cartoon form the overall approach whereby agonist binding leads to an active state which has high affinity for Nb39, thereby increasing the BRET signal, B shows a typical kinetic plot showing the effect of the agonist morphine and the antagonist naloxone. C shows a typical kinetic plot at 2 different sodium concentrations along with a no-drug control. D. Shows the response due to the agonist morphine for N=3 independent biological replicates. E shows the response due to the antagonist naloxone at two different sodium concentrations for N=3 independent biological replicates. For all: ****p<0.001

We next tested the hypothesis that sodium stabilizes inactive conformations by varying extracellular sodium concentrations and replacing depleted NaCl with choline chloride as described (*28*). As seen in Fig 3C, the apparent conformational transitions stabilized by agonist were greatly attenuated in 140 mM NaCl and 60 mM NaCl vs 0 mM NaCl. This is consistent with sodium as a potent NAM, as predicted by radioligand binding studies. Interestingly, the effect of naloxone for reversing the agonist-stabilized state was insensitive to extracellular sodium as measured by Nb6 association (Fig 2E?). In a similar manner, we evaluated the effect of extracellular sodium on modulating agonist- and antagonist-stabilized states using Nb39, which recognizes the active state of MOR. As shown (Fig 3B), the ability of morphine to induce conformational transitions to the active state was attenuated in 140 mM NaCl when compared with 0 mM NaCl. Interestingly, the basal BRET was diminished in 0 mM NaCl while the naloxone-stabilized transition was enhanced in 140 mM NaCl when compared with 0 mM NaCl (Fig 3D and 3E).

We next evaluated the bitopic MOR agonists C5- and C6-guano, which interact with both the sodium and the orthosteric sites of MOR(*27*). Both compounds are partial agonists when compared with the full agonist DAMGO. As shown in Fig 4, both C5- and C6-guano caused a smaller conformational stabilization for the active-state sensing nanobody Nb39 when compared with morphine. Importantly, neither C5-nor C6-guano’s effect on conformational dynamics at MOR were sensitive to changes in extracellular sodium concentrations.

**Fig 4.**
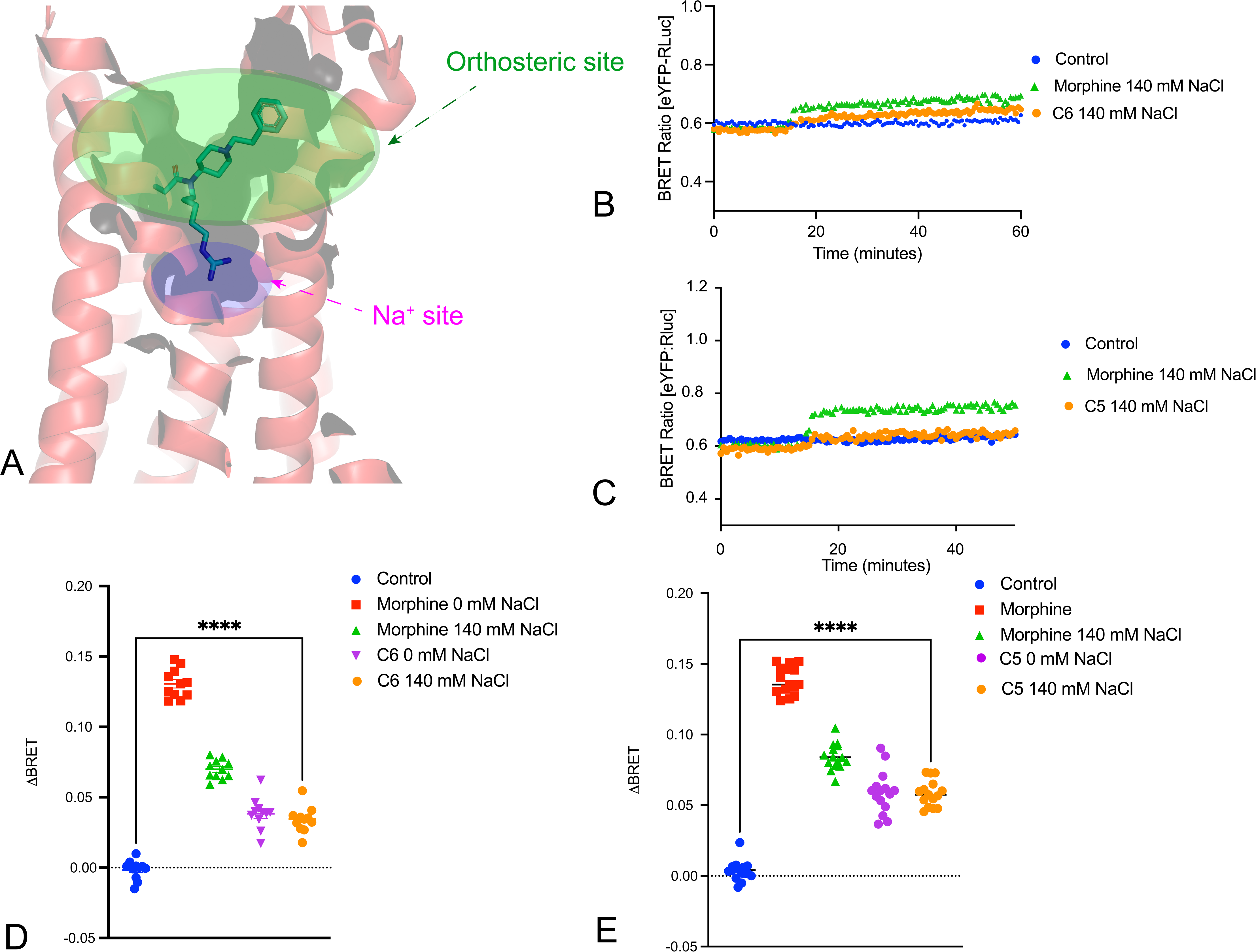
Bitopic MOR agonist-induced conformational transitions are unaffected by sodium. Panel A shows the binding pose of the bitopic ligand to MOR (PDB: 7U2K). Panels B and C show the kinetic tracings showing response of Nb39 association to morphine and C6- or C5-guano bitopic ligands. Panels D and E show the averaged BRET changes induced by various compounds. ****p<0.0001 for all.

We next examined the effects of sodium on the ability of the full agonist DAMGO to activate various Gα subunits using our TRUPATH technology (*26*). We used the NTSR1-neurotensin receptor as a control, as it has been previously shown to activate every Gα subunit except GαS (*26*) (*32*). As shown in Fig 5, the ability of neurotensin to activate GoA and G13 (and other tested Gα subunits) was insensitive to extracellular sodium concentration. When extracellular NaCl was increased to 140 mM, DAMGO’s potency at MOR decreased for all G protein pathways, while its maximal eGicacy was aGected in a transducer-dependent manner. (Fig 5; Table 2). Interestingly, the ability of DAMGO to activate Gz was enhanced in the presence of 140 mM NaCl, while DAMGO’s activation of Ggust, although minimal, was unaffected by altering extracellular sodium concentration.

**Fig 5.**
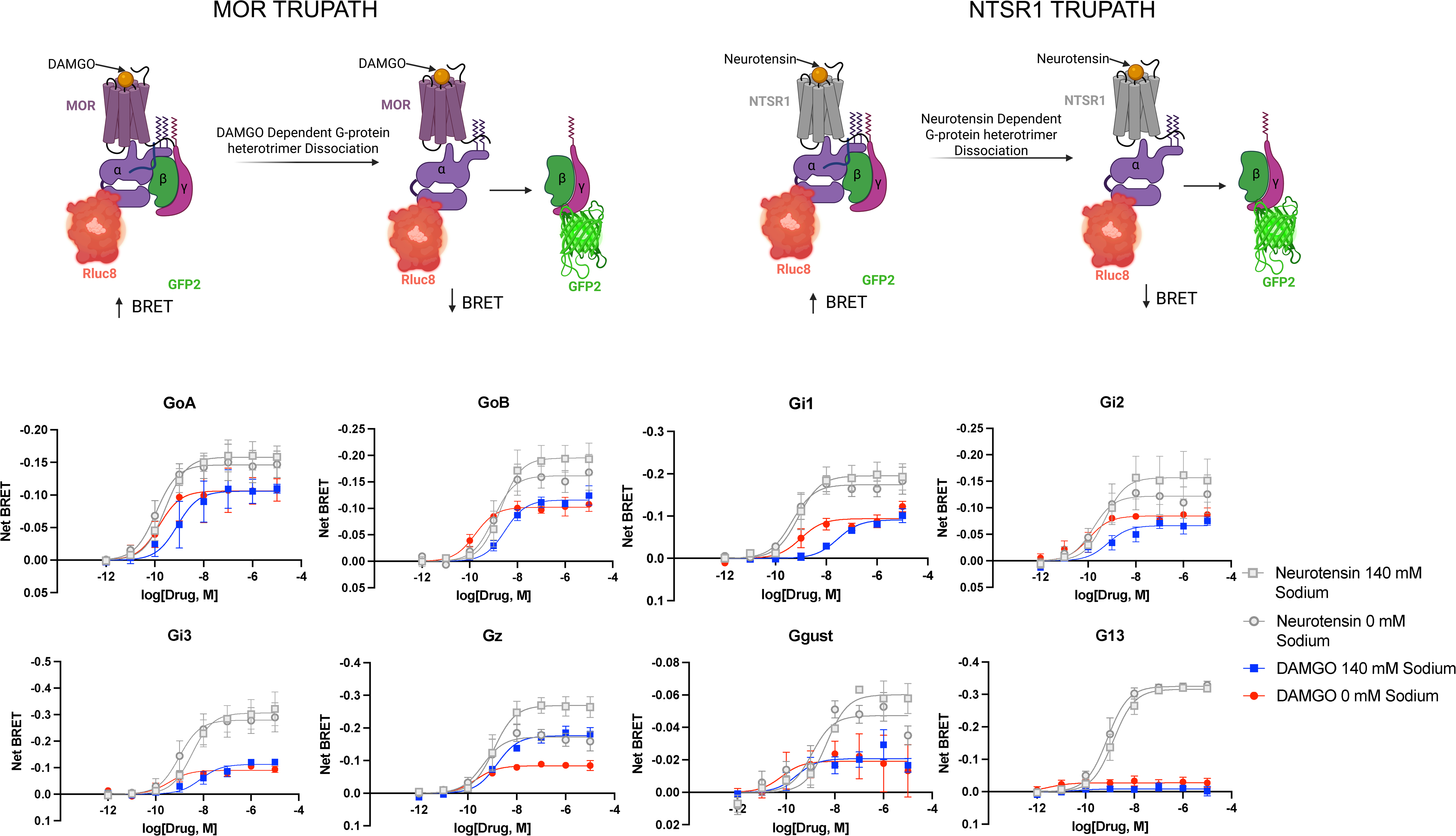
TRUPATH profiles for MOR at 0 and 140 mM NaCl. The top panel shows in cartoon form the overall strategy of the TRUPATH BRET studies. The lower two panels show the concentration-response curves for N=3 biological replicates with NTSR1 as a reference for all studies. Data represent mean +/- SEM for N=3 biological replicates.

**Table 2.** **–** Fit parameters for dose response curves on Figures 6 and 7. This data represents the mean ± SEM for N=3 biological replicates. Each section is broken out by sodium concentration, transducer, receptor, and drug. All plots are normalized to the NTSR1 response at 140 mM sodium concentration. P-values are calculated from an Extra sum-of-squares F-test between the curve fits for 140 mM Sodium and 0 mM Sodium on all parameters. ND for not detected

| Transducer | Receptor | Drug | 140 mM Sodium |  | 0 mM Sodium |  | P-value |
| --- | --- | --- | --- | --- | --- | --- | --- |
| | | | E<sub>max</sub> ± SEM | pEC<sub>50</sub> ± SEM | E<sub>max</sub> ± SEM | pEC<sub>50</sub> | |
| GoA | NTSR1 | Neurotensin | 100 ± 4.43 | 9.57 ± 0.158 | 92.6 ± 4.94 | 9.95 ± 0.188 | 0.2781 |
| | MOR | DAMGO | 67 ± 6.51 | 9.08 ± 0.312 | 67.3 ± 6.78 | 9.82 ± 0.356 | 0.3171 |
| | KOR | Salvinorin A | 101.9 ± 2.76 | 9.27 ± 0.091 | 60.6 ± 2.1 | 9.3 ± 0.118 | <0.0001 |
| | KOR | U69,593 | 107 ± 2.56 | 8.71 ± 0.075 | 58.3 ± 2.33 | 9.05 ± 0.127 | <0.0001 |
|  | KOR | Nor-BNI | ND | ND | ND | ND |  |
| GoB | NTSR1 | Neurotensin | 100 ± 6.28 | 8.71 ± 0.198 | 82.8 ± 5.55 | 9 ± 0.212 | 0.1257 |
| | MOR | DAMGO | 59.3 ± 2.55 | 8.52 ± 0.135 | 52.4 ± 2.26 | 9.77 ± 0.153 | <0.0001 |
| | KOR | Salvinorin A | 90.9 ± 3.11 | 9.86 ± 0.121 | 51.9 ± 2.28 | 9.44 ± 0.155 | <0.0001 |
| | KOR | U69,593 | 87.7 ± 3.18 | 9.05 ± 0.116 | 52 ± 2.64 | 9.31 ± 0.173 | <0.0001 |
|  | KOR | Nor-BNI | ND | ND | ND | ND |  |
| Gi1 | NTSR1 | Neurotensin | 100 ± 6.26 | 9.1 ± 0.202 | 89.2 ± 4.44 | 9.35 ± 0.172 | 0.3368 |
| | MOR | DAMGO | 46.7 ± 2.61 | 7.55 ± 0.15 | 48.1 ± 3.6 | 9.01 ± 0.237 | <0.0001 |
| | KOR | Salvinorin A | 108.3 ± 4.1 | 9.32 ± 0.129 | 77.4 ± 3.26 | 9.15 ± 0.137 | <0.0001 |
| | KOR | U69,593 | 107.3 ± 4.3 | 8.35 ± 0.122 | 75.7 ± 2.38 | 9.02 ± 0.1 | <0.0001 |
|  | KOR | Nor-BNI | ND | ND | ND | ND |  |
| Gi2 | NTSR1 | Neurotensin | 100 ± 9.09 | 9.29 ± 0.309 | 77.8 ± 7.49 | 9.8 ± 0.34 | 0.1697 |
| | MOR | DAMGO | 42.4 ± 3.63 | 9.17 ± 0.281 | 53.9 ± 2.95 | 9.97 ± 0.193 | 0.0012 |
| | KOR | Salvinorin A | 82 ± 5.91 | 9.31 ± 0.246 | 63.8 ± 2.21 | 9.14 ± 0.113 | 0.0056 |
| | KOR | U69,593 | 83.7 ± 7.15 | 8.32 ± 0.257 | 62.6 ± 3.25 | 9.1 ± 0.167 | 0.0215 |
|  | KOR | Nor-BNI | ND | ND | ND | ND |  |
| Gi3 | NTSR1 | Neurotensin | 100 ± 6.56 | 8.47 ± 0.205 | 91.1 ± 6.8 | 9.02 ± 0.237 | 0.2376 |
| | MOR | DAMGO | 36.8 ± 2.9 | 8.12 ± 0.222 | 29.4 ± 1.9 | 9.43 ± 0.226 | 0.0065 |
| | KOR | Salvinorin A | 93.3 ± 3.31 | 9.18 ± 0.116 | 73.3 ± 2.29 | 9.17 ± 0.102 | <0.0001 |
| | KOR | U69,593 | 91 ± 3.99 | 8.34 ± 0.133 | 74.1 ± 3.03 | 9.04 ± 0.13 | 0.0015 |
|  | KOR | Nor-BNI | ND | ND | ND | ND |  |
| Gz | NTSR1 | Neurotensin | 100 ± 5.05 | 8.94 ± 0.159 | 64.1 ± 4.05 | 9.38 ± 0.22 | <0.0001 |
| | MOR | DAMGO | 65.5 ± 2.43 | 8.82 ± 0.117 | 31.2 ± 1.11 | 9.66 ± 0.126 | <0.0001 |
| | KOR | Salvinorin A | 51.5 ± 1.51 | 10.45 ± 0.115 | 13.6 ± 1.27 | 9.87 ± 0.329 | <0.0001 |
| | KOR | U69,593 | 57.4 ± 1.75 | 9.89 ± 0.107 | 11.6 ± 0.99 | 9.74 ± 0 | <0.0001 |
|  | KOR | Nor-BNI | ND | ND | ND | ND |  |
| Ggust | NTSR1 | Neurotensin | 100 ± 5.86 | 8.34 ± 0.177 | 78.6 ± 7.1 | 8.95 ± 0.285 | 0.0537 |
| | MOR | DAMGO | 34.5 ± 4.24 | 9.57 ± 0.438 | 31.9 ± 7.63 | 10.2 ± 0.879 | 0.8089 |
| | KOR | Salvinorin A | 155.3 ± 4.26 | 8.52 ± 0.086 | 149.4 ± 4.26 | 8.78 ± 0.09 | 0.1939 |
| | KOR | U69,593 | 157.5 ± 7.67 | 8.09 ± 0.136 | 159.2 ± 4.75 | 8.52 ± 0.094 | 0.0364 |
| | KOR | Nor-BNI | 25.2 ± 7.07 | 6.85 ± 0.616 | 28.1 ± 4.62 | 7.59 ± 0.444 | 0.4192 |
| G13 | NTSR1 | Neurotensin | 100 ± 2.76 | 8.86 ± 0.087 | 103 ± 3 | 9.11 ± 0.094 | 0.0519 |
|  | MOR | DAMGO | ND | ND | ND | ND |  |
|  | KOR | Salvinorin A | ND | ND | ND | ND |  |
|  | KOR | U69,593 | ND | ND | ND | ND |  |
|  | KOR | Nor-BNI | ND | ND | ND | ND |  |

### Real-time visualization of sodium-stabilized conformational transitions at KOR

We next examined the κ-opioid receptor (KOR), as prior studies have indicated that KOR agonist binding is relatively insensitive to extracellular sodium concentrations (*33*) (*34*) (*35*). In initial saturation binding studies, we found a decrease in agonist affinity (K_D_) and maximum number of binding sites (B_MAX_) comparing results obtained in buffer containing 140 mM NaCl vs 0 mM NaCl (Fig 6B-C). Significantly, the agonist-induced conformational transitions were insensitive to extracellular sodium (Fig 6D-F). When comparing the ability of the full agonists salvinorin A (*3*) and U69,593(*36*) to facilitate KOR activation of hetereotrimeric G proteins we again used our TRUPATH resource (*26*). Surprisingly, we found that the responses to salvinorin A and U69,593 were potentiated in 140 mM NaCl vs experiments performed in 0 mM NaCl, with no effect on neurotensin’s activity at NTSR1 (Fig 7). We also observed that the inverse agonist activity of Nor-BNI was transducer and sodium-dependent at GoA and GoB, and minimal detectible inverse agonist activity at other tested G proteins (Fig 7).

**Fig 6.**
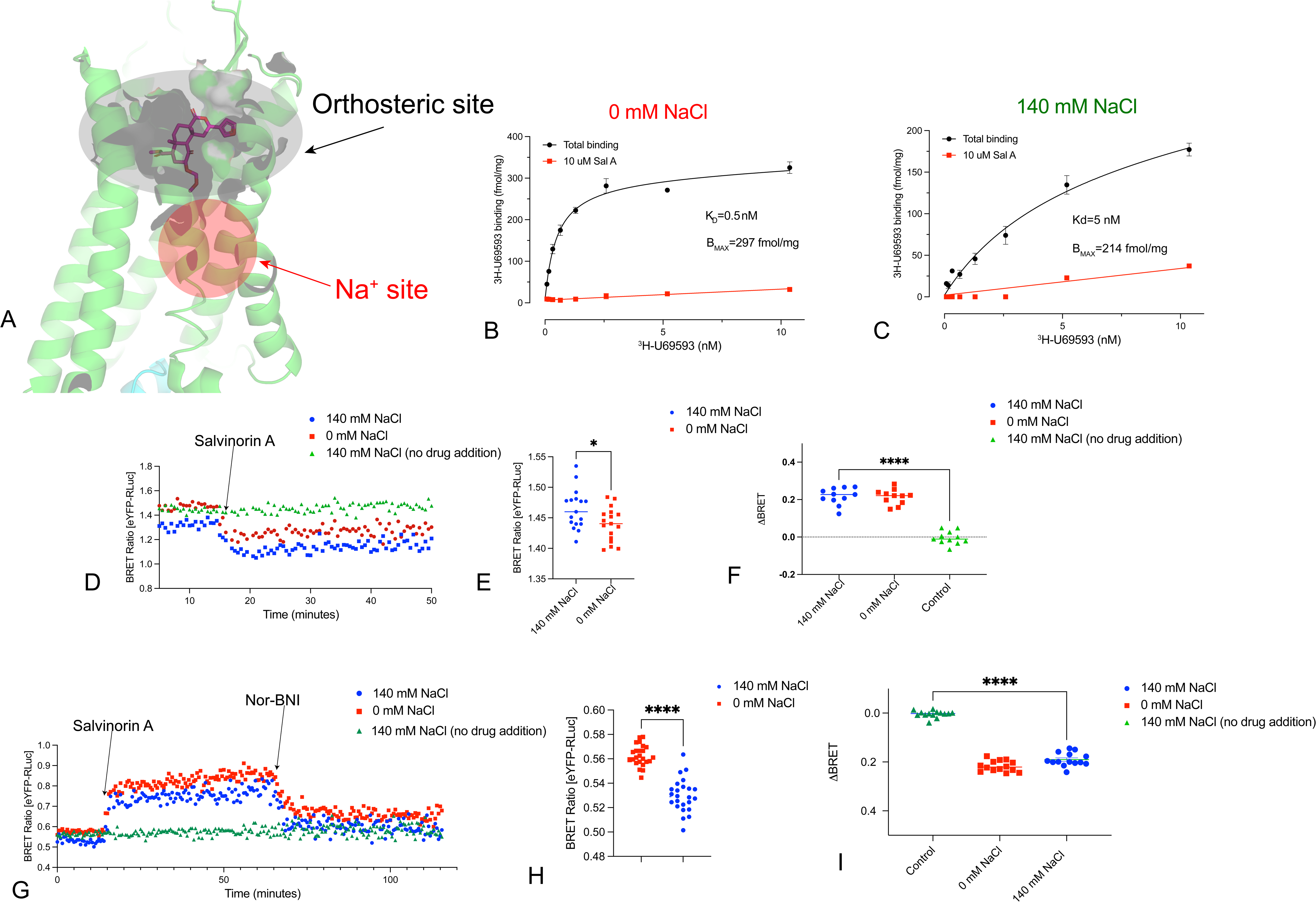
Real-Time Conformational Measurement (RTCM) BRET at KOR using the inactive- and active-state nanobodies. Panel A shows the location of the orthosteric and allosteric sodium sites in KOR. Panels B and C shows saturation binding isotherms for KOR at 0 and 140 mM NaCl, respectively. Panel D shows a kinetic plot of KOR using Nb39 as reporter while Panels E and F show the averaged response for N=3 biological replicates. Panel G shows a kinetic plot of KOR using Nb6 as reporter while Panels H and I show the averaged response for N=3 biological replicates. ****p<0.0001 for all.

**Fig 7.**
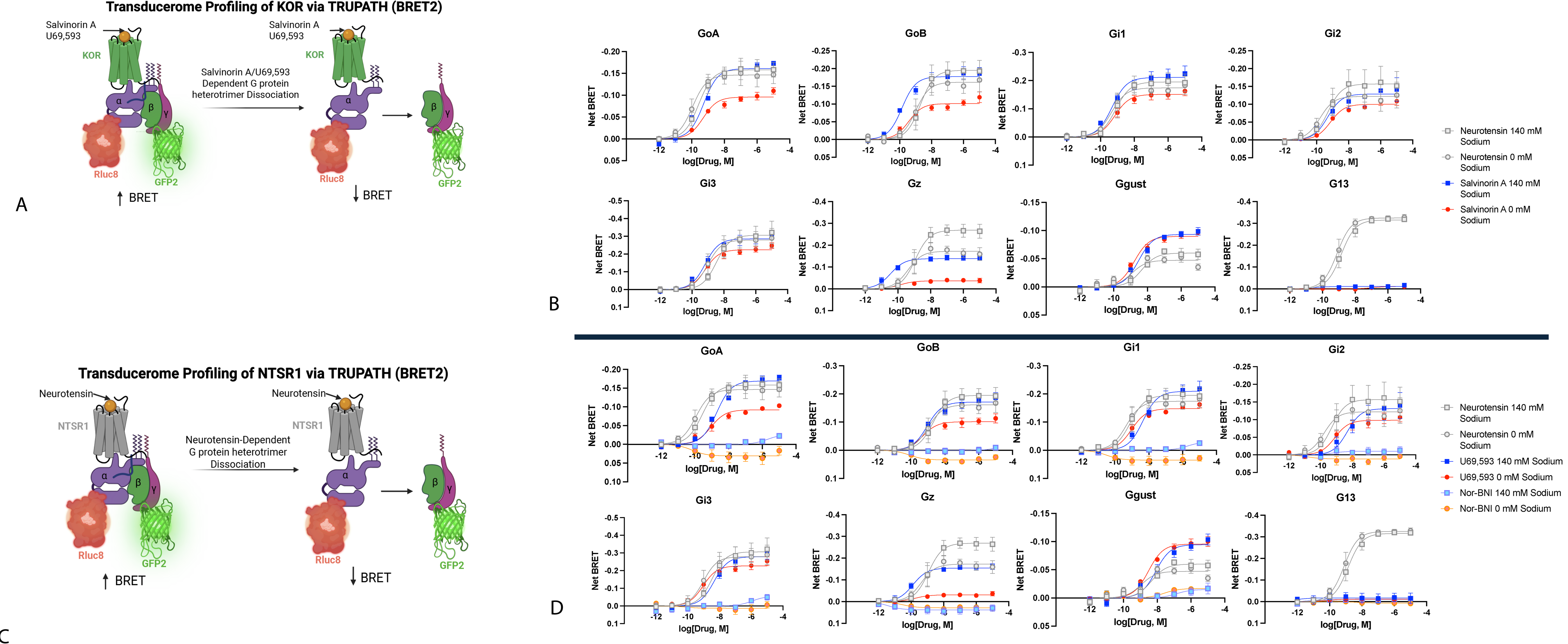
TRUPATH profiles for KOR at 0 and 140 mM NaCl. The left panels show in cartoon form the overall strategy of the TRUPATH BRET studies. The right two panels show the concentration-response curves for N=3 biological replicates with NTSR1 as a reference for all studies. Data represent mean +/- SEM for N=3 biological replicates.

## DISCUSSION

The main finding of this paper is that sodium’s allosteric modulation of GPCRs is complex with differential and opposing actions on ligand affinities, agonist- and antagonist-induced conformational transitions and receptor-transducer coupling. At first glance, these findings are counter-intuitive given the large number of prior radioligand binding studies which have demonstrated sodium’s negative allosteric effect on agonist binding affinity (for review see ref (*24*)). We note, however, that several prior studies have examined the effects of extracellular sodium on GPCR signaling *in vitro* with results reminiscent of ours. Thus, either inhibition, potentiation or no effect of sodium has been reported for many GPCR signaling systems. For the ghrelin receptor reconstituted with G proteins, sodium inhibits agonist-stimulated GTP hydrolysis (*37*). Similarly, for the H_3_-histamine(*38*). DOR(*39*), Y2- and Y4-Neuropeptide Y, Formyl-peptide (*40*) receptors agonist-stimulated signaling was inhibited by sodium ions. By contrast, no effect of sodium was seen for H_4_-histamine(*41*), CB_1_-and CB_2_-cannibinoid receptors (*42*), while stimulation by sodium has been reported for the β_2_-adrenergic (*43*), the fMLP(*44*), CXCR4-chemokine (*45*) receptors. As is clear from our results with various Gα subunits, there are differential and significant effects of sodium on coupling of distinct GPCRs to distinct transducers; these findings are, to our knowledge, new.

Given these divergent effects of sodium, what is the expected effect on signaling in vivo by these various receptors? As shown in Fig 8, at the cell surface GPCRs will be exposed to relatively high sodium concentrations (∼140 mM), while the intracellular concentration is low (7-10 mM) and the concentration within the endosomal compartment being intermediate (70-100 mM). Prior results showed that intracellular and extracellular opioid receptors, when isolated, have equivalent sodium sensitivities (*46*) (*47*). Thus, one can predict that intracellular μ opioid receptor-G-protein complexes, for instance, would have relatively higher activities and higher potencies for cell-permeable opioids. By contrast, our results predict that cell-permeable KOR agonists will show greater potencies and efficacies for cell surface as compared with intracellular receptors. For NTSR1, minimal effects of sodium were observed, suggesting that intracellular and cell surface NTRS1 receptors will show similar coupling efficiencies to G proteins.

**Fig 8.**
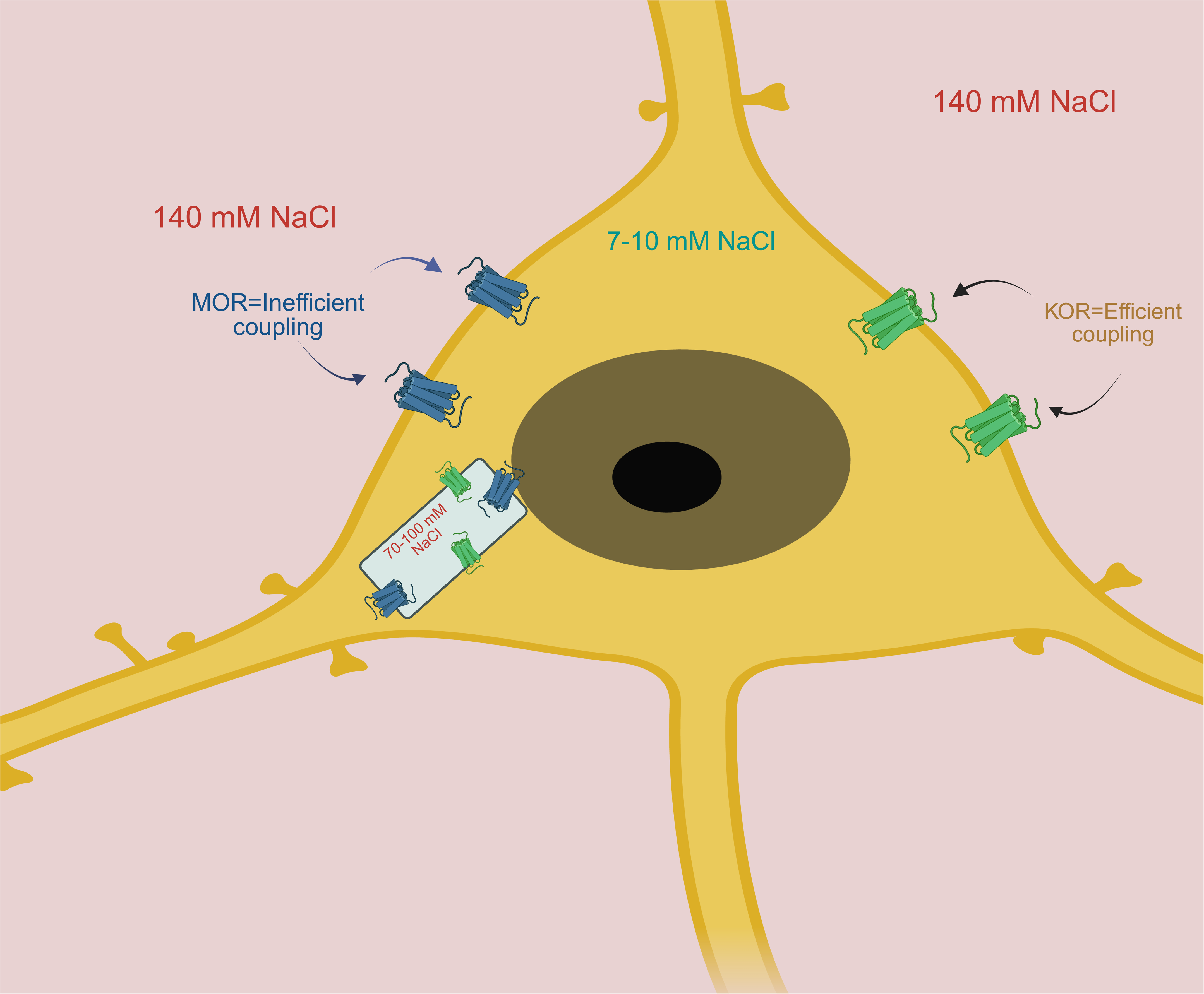
Cartoon depicting effects of varying sodium at distinct subcellular compartments on MOR and KOR signaling. As can be seen, cell surface MOR are predicted to have lower potencies and efficacies for agonists compared with KOR, while intracellular receptors will display intermediate effects of sodium.

Taken together, these findings imply that the actions of sodium will vary depending upon the GPCR, its subcellular localization, and the nature of the transducers engaged. Bitopic ligands that interact with both the sodium and orthosteric site, as shown here, remain relatively unaffected by sodium and, hence, their actions on intracellular and cell-surface receptors will be equivalent. Given the diversity of GPCR-transducer combinations, and the wide variation in intra-vs extracellular sodium concentrations under physiological conditions, our results imply a heretofore unappreciated complexity encoded by the interaction of a single ion with similar GPCRs.

## Acknowledgements

This work was supported by the NIMH Psychoactive Drug Screening Program, U19NS138975, R37DA045657 and RO1MH112205 to BLR, R01DA059978 and R01DA057790 to S.M..

---

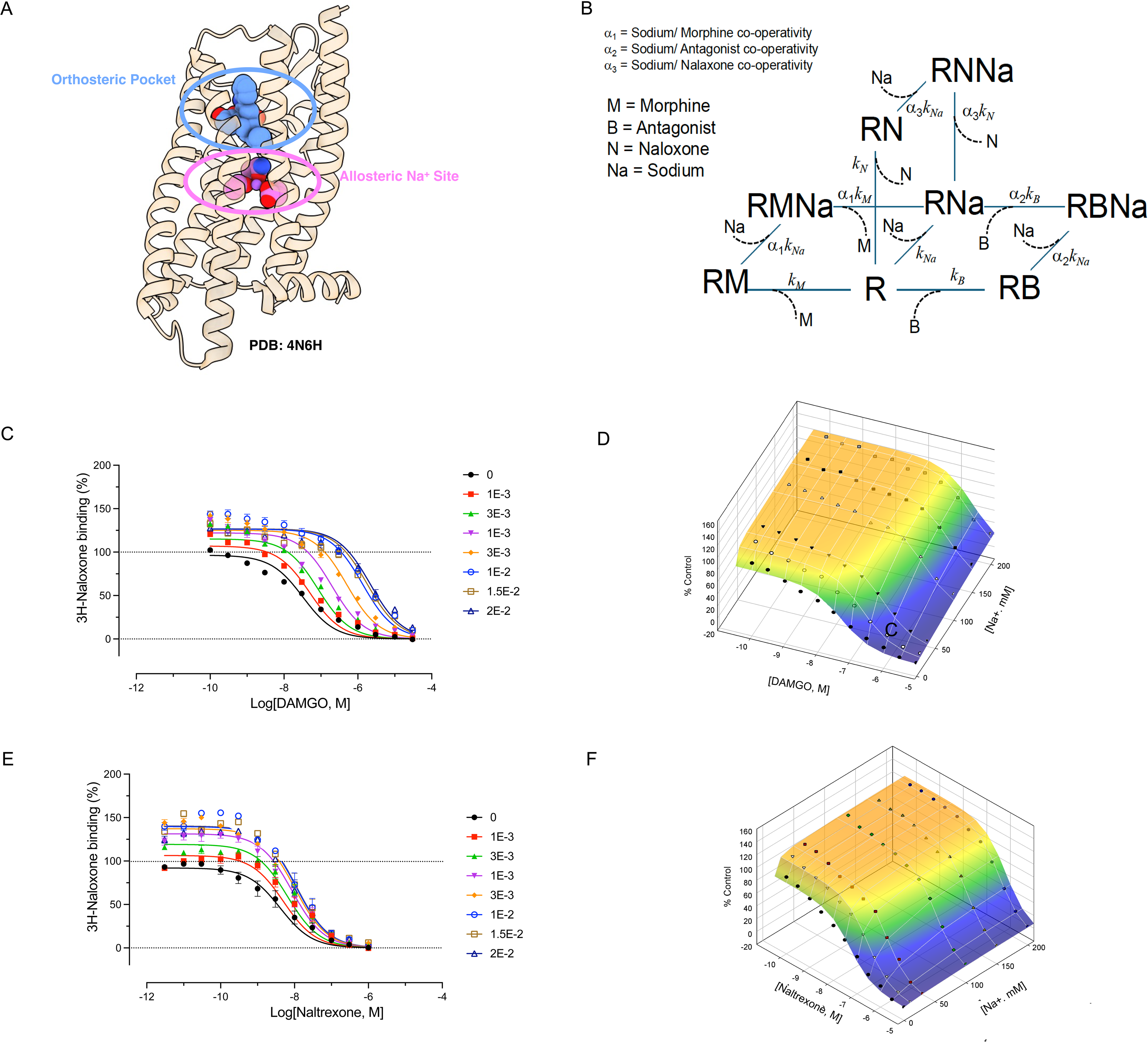

